# Reduced prevalence of phage defense systems in *Pseudomonas aeruginosa* strains from Cystic Fibrosis Patients

**DOI:** 10.1101/2024.11.16.623971

**Authors:** Daan F. van den Berg, Stan J.J. Brouns

## Abstract

Cystic fibrosis is a genetic disorder that affects mucus clearance, particularly of the lungs. As a result, cystic fibrosis patients often experience infections from bacteria, which contribute to the disease progression. *Pseudomonas aeruginosa* is one of the most common opportunistic pathogens associated with cystic fibrosis. The presence of *P. aeruginosa* complicates the treatment due to its high antibiotic resistance. Thus, research is ongoing to treat these infections with bacterial viruses instead, known as bacteriophages. Notably, *P. aeruginosa* clinical strains possess a variety of phage defense mechanisms that may limit the effectiveness of phage therapy. In this study, we compared the defense system repertoire of *P. aeruginosa* strains isolated from cystic fibrosis patients with those from non-cystic fibrosis patients. Our findings reveal that *P. aeruginosa* strains isolated from cystic fibrosis patients have fewer phage defense mechanisms per strain than from non-cystic fibrosis patients, suggesting altered phage selection pressures in strains colonizing CF patient lungs.

**Importance:** Cystic fibrosis patients often experience chronic *Pseudomonas aeruginosa* lung infections, which are challenging to treat with antibiotics and contribute to disease progression and eventual respiratory failure. Phage therapy is being explored as an alternative treatment strategy for these infections. However, assessing strain susceptibility to phage treatment is essential for ensuring efficacy. To address this, we investigated whether CF-associated clinical *P. aeruginosa* strains have a distinct phage defense repertoire compared to those isolated from other lung patients. We observed that CF-associated *P. aeruginosa* strains have significantly fewer phage defenses, possibly affecting the susceptibility of these strains to phage infection.

## Introduction

Cystic fibrosis (CF) is a genetic disorder that affects mucus clearance, especially of the lungs^1^. The decrease in mucus clearance creates a breeding ground for opportunistic pathogens, such as bacteria, to colonize the lungs^1^. These infections are often chronic, accumulate resistance to antibiotic treatment, and cause major lung damage^1,2^. Over time, this damage results in respiratory failure, the most common cause of death for CF patients^1^. Lung infections in CF patients are often caused by strains of *Pseudomonas aeruginosa*^1^. Although antibiotic treatments can alleviate patient symptoms and prolong life, they generally fail to fully eradicate *P. aeruginosa* from the lungs^3^. Consequently, research has turned to alternative treatment options, including the use of bacterial viruses (phages)^4,5,6,7,8^. However, the efficacy of phage-based treatments might also be limited since *P. aeruginosa* strains often possess a variety of phage defense mechanisms that may limit the sensitivity of strains to phages^9^. Particularly, a recent study on the abundance of defense systems in clinical isolates of *P. aeruginosa* found that these strains accumulate a large number of defense systems and are less susceptible to phage infection^9^. However, it is unknown if the distribution of phage defense systems varies with the pathogen environment or patient conditions. In this study, we investigated the defense system repertoire of clinical *P. aeruginosa* isolates from mucus-rich lungs of CF patients by comparing them to strains isolated from non-CF lung patients.

## Results and discussion

We utilized the Pseudomonas Genome Database to investigate the defense system repertoire of *P. aeruginosa* isolates from the respiratory system of CF and non-CF (e.g. pneumonia) patients^10^. We first compared the prevalence of the phage defense systems and found isolates from CF patients to encode significantly fewer phage defense systems per strain (non-CF: 9.8 vs CF: 6.8; a 30% reduction; Welch Two Sample t-test, t = −9.878, df = 601.37, p-value < 2.2^−16^) (Figure 1A). Interestingly, this reduction in the number of defense systems per strain did not affect the overall diversity of the defense repertoire, where the Shannon index (Both: 3.86) and evenness (Both: 0.80) remained identical (Table S1). Therefore, we hypothesized that the decrease in defense systems per strain is not the result of a broad, non-specific reduction. Instead, we suspected that the overall reduction is possibly caused by differences in the abundance of a few prevalent defense systems. To assess this, we compared the abundance of each defense system in CF vs non-CF lung isolates (Table S1), considering changes greater than 2-fold (log_2_FC > 1 or log_2_FC < −1) as relevant (Table S2). We identified 31 defense systems with considerable changes in abundance (seven were more abundant and 24 were less abundant in CF strains; Figure 1B-C, SC-E), but only four of these systems were prevalent enough to potentially affect the overall number of defense systems per strain (present in more than 10%) in either group, all four were more depleted in CF isolates. These four defense systems include Wadjet type I^11^ (non-CF: 37.8% vs CF:13.4%; log_2_FC = −1.4), Shedu^12^ (non-CF: 23.5% vs CF: 9.7%; log_2_FC = −1.2), RM type III (non-CF: 23.5% vs CF: 7.7%; log_2_FC = −1.5) and Zorya type I^13^ (non-CF: 24.4% vs CF: 3.4%; log_2_FC = −2.5). The reduced prevalence of just these four defense systems accounts for almost half the reduction in defense system number per strain observed in CF compared to non-CF isolates. Besides these defense systems, CRISPR-Cas type IV-A is also noteworthy for being completely absent in the CF isolates, while present in the non-CF isolates (non-CF: 2.6% vs CF: 0%; log_2_FC = −1.8). Interestingly, all of the above-mentioned defense systems act on foreign DNA, and notably, two of these defense systems, Wadjet type I and CRISPR-Cas type IV-A, are known to act upon plasmids specifically^11,14,15^. The reduced presence of plasmid-restrictive mechanisms of these strains may highlight the role of plasmids in CF-infecting strains, potentially facilitating their acquisition to gain resistance against antibiotics used for treating CF infections^16,17^.

**Figure 1.**
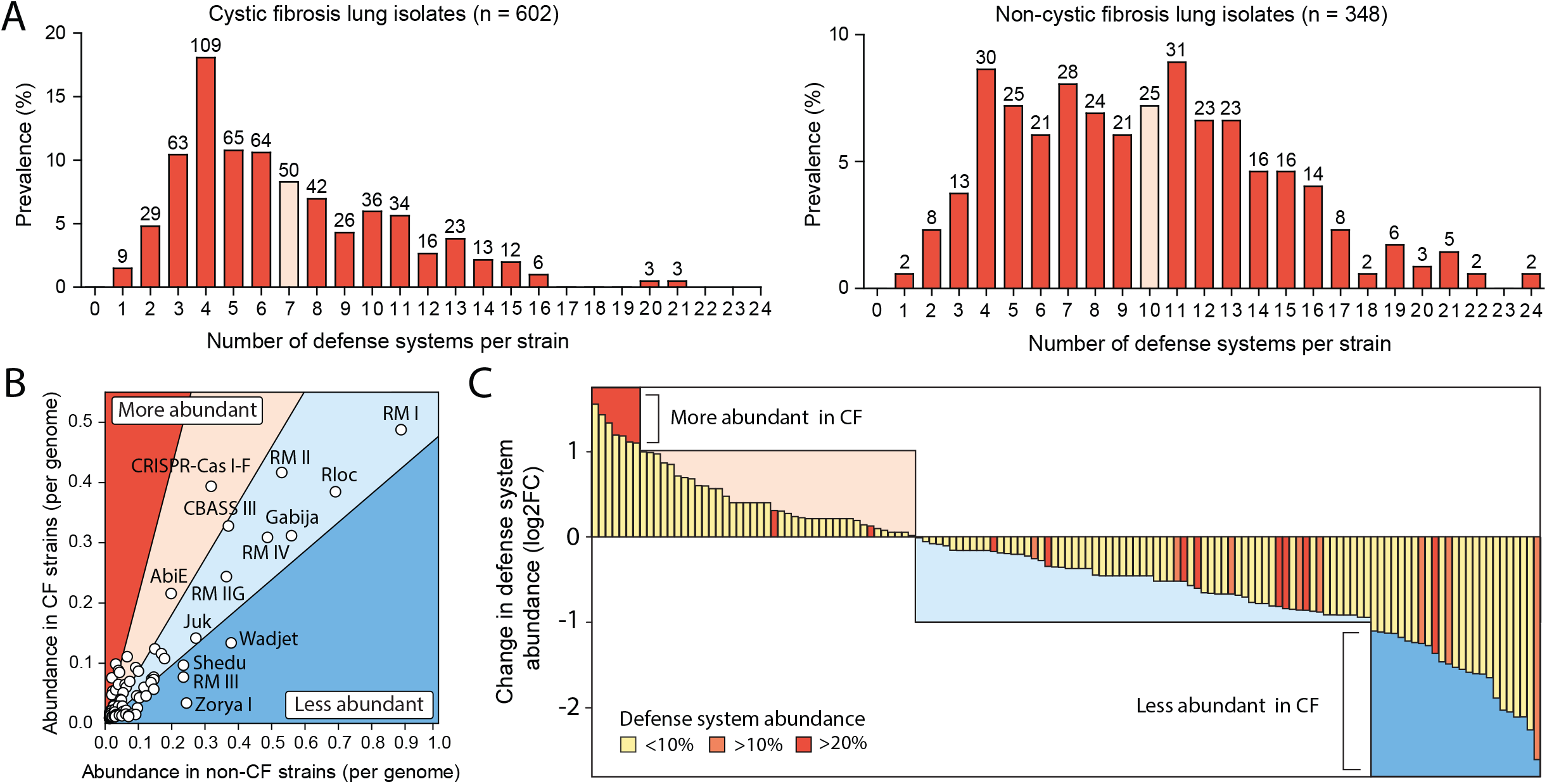
*Pseudomonas aeruginosa* strains isolated from cystic fibrosis (CF) patient lungs have fewer defense systems. (A) The number of defense systems per *P. aeruginosa* strain isolated from CF lungs (left) and non-CF lungs (right). The median number of defense systems per strain is indicated with a lighter red colored bar. (B) The comparative abundance of individual defense systems in *P. aeruginosa* strains isolated from CF and non-CF lungs. The graph is divided into four sections: Red represents a log_2_FC greater than 1, light red indicates a log_2_FC between 0 and 1, light blue corresponds to a log_2_FC between 0 and −1, and blue signifies a log_2_FC less than −1. (C) The relative log_2_FC change in the prevalence of individual defense systems in *P. aeruginosa* strains isolated from CF lungs compared to non-CF lungs. A larger depiction of this graph, which also shows the individual defense system names, can be seen in supplementary figure S1C.

It remains unclear whether the reduced prevalence of defense systems in CF isolates was caused by strain selection before, or during lung infection in CF patients. As for antibiotic resistance, strains infecting CF lungs adapt over time^1^, and could therefore also adapt in the context of phages and other mobile genetic elements such as plasmids. In the CF lung environment, this adaptation seems to involve the loss of phage defense mechanisms, possibly because CF lungs are less penetrable to phages due to factors such as reduced air circulation, a thicker and dehydrated mucus layer, and a higher prevalence of biofilms^1,18^. Supporting this hypothesis, we observed fewer prophages in the genomes of CF lung isolates, suggesting reduced exposure to phage predation (CF: 3.3 vs non-CF: 5.3 prophages per genome; 38% decrease; Welch Two Sample t-test: t = −12.379, df = 508.88, p-value < 2.2^−16^) (Figure S1F). Besides the reduced phage defenses of *P. aeruginosa* isolates from CF lung isolates we also observed these strains to have a significantly smaller genome (CF: 6.5 Mb vs non-CF:

6.8 Mb; a 4% decrease; Welch Two Sample t-test: t = −13.811, df = 594.47, p-value < 2.2^−16^) and to encode fewer genes (CF: 6053 vs non-CF: 6298 genes; a 4% decrease; Welch Two Sample t-test: t = −11.715, df = 625.07, p-value < 2.2^−16^) (Figure S1G,H). These characteristics reflect a more adapted and specialized genome of *Pseudomonas aeruginosa* strains in the CF lung environment^19,20,21^.

## Conclusion

We observed that *P. aeruginosa* strains isolated from the lungs of CF patients encode a more limited number of phage defense systems compared to strains isolated from non-CF patient lungs. This can largely be attributed to a reduced abundance of several key phage defense systems including Wadjet, Shedu, Zorya, and RM type, which likely results from adaptation to the CF lung environment. This provides a promising perspective that these bacterial strains are more susceptible to phage therapeutic options.

## Method

All complete *P. aeruginosa* genomes were downloaded from the Pseudomonas Genome Database on the 9th of April 2023 (n = 14,230). The metadata was utilized to select the strains for: Species = *Pseudomonas aeruginosa*, Assembly version status = latest, Host taxonomic name = Homo sapiens. A further selection was then made between isolates from the respiratory system of CF (n = 602) and those of non-CF (n = 348) (Table S1). The non-CF patients primarily were affected by unspecified respiratory tract infection (57.5%), ventilator-associated pneumonia (20.1%), and pneumonia (12.4%). Chronic bronchitis and chronic obstructive pulmonary disease were less frequently prevalent, 4.9% and 0.3% respectively. Defense systems were detected in *P. aeruginosa* genomes with DefenseFinder v1.3.0^22^ using DefenseFinderModels v1.3.0. Prophages were identified using PhiSpy v4.2.21 (default settings)^23^.

## Acknowledgments

We thank members of the Brounslab for the many discussions and ideas that improved our work. Especially, dr. Nadiia Pozhydaieva and dr. Ana Rita Costa for critically reading and providing feedback on the manuscript.

---

**Figure S1. Comparison of defense system repertoire in *Pseudomonas aeruginosa* strains isolated from cystic and non-cystic fibrosis lungs**. (A) The prevalence of individual defense systems per *P. aeruginosa* strain isolated from CF lungs. (B) The prevalence of individual defense systems per *P. aeruginosa* strain isolated from non-CF lungs. (C) The relative log_2_FC change in prevalence of individual defense systems between *P. aeruginosa* strains isolated from CF and non-CF patients. (D) The subset of defense systems that are found to be overly abundant (log_2_FC > 1) and (E) less abundant (log_2_FC < −1) in CF isolates. (F) The number of detectable prophages in *P. aeruginosa* strains isolated from CF and non-CF patients. (G) The genome size of *P. aeruginosa* strains isolated from CF and non-CF patients. (H) The number of genes present in *P. aeruginosa* strains isolated from CF and non-CF patients.

**Table S1**. Summary table of the *Pseudomonas aeruginosa* strains and additional information, including the defense system repertoire.

**Table S2**. The relative fold changes in defense system prevalence in *Pseudomonas aeruginosa* strains isolated from CF lungs compared to those from non-CF lungs

